# Enhancing Prediction of Human Traits and Behaviors through Ensemble Learning of Traditional and Novel Resting-State fMRI Connectivity Analyses

**DOI:** 10.1101/2024.03.30.587447

**Authors:** Takaaki Yoshimoto, Kai Tokunaga, Junichi Chikazoe

## Abstract

Recent efforts in cognitive neuroscience have focused on leveraging resting-state functional connectivity (RSFC) data from fMRI to predict human traits and behaviors more accurately. Traditional methods typically analyze RSFC by correlating averaged time-series data between regions of interest (ROIs) or networks, a process that may overlook critical spatial signal patterns. To address this limitation, we introduced a novel linear regression technique that estimates RSFC by predicting spatial brain activity patterns in a target ROI from those in a seed ROI. We applied both traditional and our novel RSFC estimation methods to a large-scale dataset from the Human Connectome Project, analyzing resting-state fMRI data to predict sex, age, personality traits, and psychological task performance. Additionally, we developed an ensemble learner that integrates these methods using a weighted average approach to enhance prediction accuracy. Our findings revealed that hierarchical clustering of RSFC patterns using our novel method displays distinct whole-brain grouping patterns compared to the traditional approach. Importantly, the ensemble model outperformed the traditional RSFC method in predicting human traits and behaviors. Notably, the predictions from the traditional and novel methods showed relatively low similarity, indicating that our novel approach captures unique and previously undetected information about human traits and behaviors through fine-grained local spatial patterns of neural activation. These results highlight the potential of combining traditional and innovative RSFC analysis techniques to enrich our understanding of the neural basis of human traits and behaviors.

## Introduction

One of the major goals of systems neuroscience is to elucidate brain-behavior relationships. Resting state functional connectivity (RSFC) is the spontaneous synchronization of brain activity between different brain regions in the absence of explicit tasks (Biswal et al., 1995). RSFC measured by fMRI technique has been found to contain information about human sex differences (Fan et al., 2020, Zhang et al., 2018), age (Geerligs et al., 2015, Varangis et al., 2019, Kim et al., 2022), personality tendencies (Hsu et al., 2018 Gentili et al., 2017), intelligence (Fan et al., 2020, Hearne et al., 2016), performance on psychological tasks (Liégeois., 2019, Li et al., 2019, He et al., 2020) and classification of neuropsychiatric disorders (Yamashita et al. 2020, Yahata et al., 2016, Rashid et al., 2016) in studies over the last decade. As a result, there have been many attempts to develop a machine learning model with superior prediction performance for behavioral traits and diseases using RSFC data.

The development of these models involves the creation of complex models with deep neural networks (Fan et al., 2020, Zhang et al., 2018, He et al., 2020), the application of graph theory join indices (Varangis et al., 2019, Kim et al. 2022), and various other analysis algorithms, but in most cases, the correlation of averaged time-series data between ROIs or networks is used as a measure of RSFC. Although this method is computationally easy and has contributed to the development of various current training algorithms (Bijsterbosch et al., 2020), it compresses and loses information on spatial signal patterns in local areas acquired at high resolution, which is one of the strengths of fMRI. However, MVPA (multi-voxel pattern analysis) studies of human fMRI have repeatedly shown that there is rich information in local spatial signal patterns. For example, a fine-grained representation of the directional selectivity of visual targets has been found in the primary visual cortex (Haynes et Rees., 2005, Kamitani et Tong., 2005). Similarly, MVPA studies suggested that there are areas in the gustatory cortex that respond to basic tastes (Chikazoe et al., 2019). Heinzle et al. also developed a method to estimate the signal in V3 from the signal pattern in the local space of V1 and found functional connectivity which has a different nature than the conventional method of averaging local signals (Heinzle et al., 2011). These results suggest that by utilizing the rich information contained in local spatial signal patterns, it is possible to find RSFC which has a novel nature.

The novel RSFC found in this way are thought to contain information about brain-behavior relationships of a different nature than those found by traditional methods using simple averages within ROIs. By integrating information of these different natures, it is quite possible to develop a learner with better predictive performance for individual traits. In the field of machine learning, a method called ensemble learning, in which multiple weak learners are integrated to create a stronger learner with better performance, has been established. The most direct way to improve the performance of ensemble learners is to input features with qualitatively different information to each weak learner (Zhou, 2012). For example, different whole brain atlases can be used for different weak learners. By extracting multiple patterns of functional connectivity from human resting-state fMRI data using ROIs based on different whole brain atlas and different parcellation algorithms to create individual learners, and then integrating these with ensemble learning methods, ensemble leaners achieve better performance in age prediction tasks (Khosla et al., 2019) than single ones. Thus, it is quite possible that an ensemble learner of models constructed by different features based on patterns of different RSFCs will outperform a learner from a single RSFC.

In this study, we developed a novel method for estimating resting-state functional connectivity by predicting the spatial brain activity pattern in the target ROI from the spatial brain activity pattern in the seed ROI using a linear regression method and calculating the correlation between the estimated brain activity pattern in the target ROI and the actual one. We applied the traditional and novel methods to the Human connectome project’s (HCP) large-scale human resting-state fMRI data and used a support vector machine algorithm to predict sex, age, personality traits, and psychological task performance by whole-brain RSFC. Furthermore, we created an ensemble learner by combining these two learners using the weighted average method and performed the same prediction.

## 2. Materials and Methods

### 2.1. Datasets

We used the S1200 release of resting-state fMRI distributed by the Human Connectome Project (HCP; http://humanconnectomeproject.org/) (Van Essen et al., 2013). The fMRI data consisted of two days and two different phase encoding directions, a total of 4 runs. The repetition time (TR) is 0.72 s. Although the total number of participants in the project is over a thousand, we used only 995 participants who completed all 4 runs. We used rs-fMRI scans processed using the ICA+FIX pipeline with MSMAll registration (Glasser et al., 2013). The data were band-pass filtered for frequencies between 0.01 and 0.1 Hz. The original data consisted of 1,200 time-frames in each run, but we trimmed the first and last 50 frames due to signal instability and artifact caused by temporal filtering. Nuisance regressors were regressed out from the time course for each vertex. These regressors consisted of averaged white matter signal, averaged ventricular signal and global signal.

### 2.2 Extraction of two types of functional connectivity (FC)

Recently, most resting-state functional connectivity studies have used the correlation between averaged time series of two different ROIs or two different networks as a measure of connectivity strength. Hereafter, we refer to this method as the “traditional method”. In contrast, we developed a method to extract resting-state FCs by using fine-grained local spatial activation patterns. We refer to this method as “novel method”. Similarly, we refer to the FC patterns generated by these methods as “traditional FC” and “novel FC”. In both methods, we used 180 gray matter regions of interest (ROIs) in each hemisphere defined by Glasser’s scheme (Glasser et al., 2016). Note that the procedures of both methods were repeated for each subject and for each run. In this section, the main processing was performed by in-house Matlab R2019b codes.

#### 2.2.1 Traditional method

We averaged preprocessed time courses of vertices within the ROI. Using a sliding window approach (Qin et al., 2015), the FC of each ROI pair was computed using Pearson correlation coefficients with a temporal window size of 23.04 s (32 TRs). All FCs were then averaged for each connectivity.

#### 2.2.2 Novel method

Fig. 1A shows the basic procedure of this method. Basically, the connectivity strength was measured by estimating how well the neural activation pattern of the target ROI for each time point was predicted by the best-fitting linear summation of the neural activations of the seed ROI vertices. Prior to the main procedure, the time series were standardized using a Z-score transform. Each run’s data was divided into 5 folds, of which 4 folds were used to estimate regression coefficients and the remaining fold was used to estimate connectivity, as shown in Fig. 1A. To illustrate the concept of the method, both the number of time points and the number of target vertices have been reduced to three in this figure. The ridge regression technique was used to estimate the regression coefficients, as shown in Fig. 1A left. The ridge parameter was fixed at 0.1, which was predetermined. Then sum of the products of each regression coefficient and neural activation in each target vertex was computed for each time point as shown in Fig 1A right. The procedure generated predicted neural activation in the whole brain vertices at every time point. The Pearson correlation coefficient between predicted and actual spatial activation patterns in the target ROI was computed as a functional connectivity metric, as shown in Fig. 1A right. Finally, these FC metrics were averaged across time points and 5 folds for a connectivity.

**Fig. 1.**
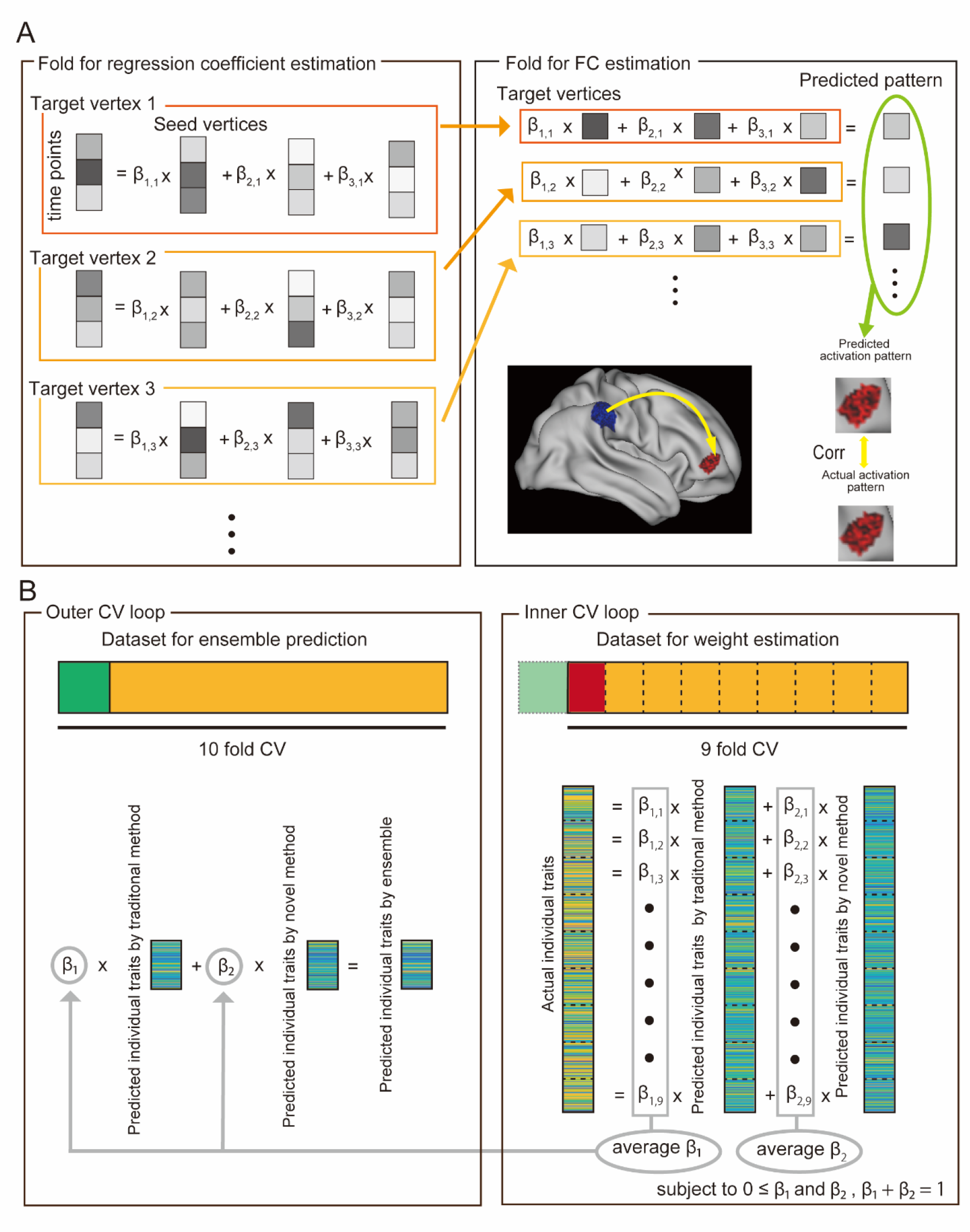
Schematic representation of the novel functional connectivity (FC) method (A) and the ensemble approach combining traditional and novel FCs (B). **A:** A conceptual diagram illustrating the procedure for the novel method. Data are partitioned into folds for the estimation of regression coefficients, as depicted on the left, with the remaining fold used to assess connectivity, displayed on the right, using a cross-validation strategy. In the schematic, the black-and-white squares symbolize neural activation at each vertex. The notation for the regression coefficients (betas) includes two parts: the first part indicates the seed vertices, while the second part refers to the target vertices. For simplicity and clearer comprehension, both the number of time points and the number of target vertices are reduced to three, differing from the complexity of actual data. This approach allows for the derivation of a connectivity metric at every time point, offering a detailed contrast to the broader estimates provided by many traditional functional connectivity (FC) analysis methods. In the diagram’s left section, encased in an orange rectangle, a regression process estimates regression coefficients for each target vertex from all seed vertices. Conversely, on the figure’s right, another orange rectangle illustrates how the neural activation for each target vertex is predicted by aggregating the products of the regression coefficients and the neural activations of the seed vertices. Specifically, the neural activation of a target vertex in one fold is predicted using the seed ROI’s activation patterns and regression coefficients derived from data across four folds. This iterative method forecasts neural activations across all brain vertices for each time period. The functional connectivity metric is determined by the Pearson correlation coefficient between the predicted and actual spatial activation patterns in the target ROI, with these metrics averaged over time points and folds for a comprehensive connectivity measure. A directional arrow in the lower right brain image indicates the flow from the seed ROI (blue) to the target ROI (red). **B:** A conceptual diagram illustrating the ensemble method combining traditional and novel FCs, where the prediction target is referred to as a trait. To prevent data leakage, a nested cross-validation technique is utilized. The dataset is segmented into ten folds for the primary cross-validation loop, with nine folds designated for outer training data and the remaining fold for outer test data. The nine-fold outer training data is further divided into eight folds for inner training data and one fold for inner validation data, with this step repeated for each fold. Within this setup, predicted traits are calculated for each fold using both FC estimation methods. Subsequently, two weights are determined through the least squares method to optimize the weighted average of the two predicted traits as close as possible to the actual trait. These weights, calculated without using the outer test dataset, are then averaged across folds to enhance prediction accuracy. In the final step, the outer test data’s predicted traits are combined using the averaged weights, culminating in the ensemble predicted trait. This method emphasizes the synergy between traditional and novel FC analyses to refine trait prediction using functional connectivity and cross-validation.

### 2.3 Hierarchical clustering on FC patterns

To explore the difference between two FC patterns, we performed hierarchical clustering on these patterns. In this section, we used Python-based codes on Jupyter-Notebook. Specifically, Nilearn toolbox was used to visualize brain surface maps. First, All FC matrices in each method were averaged across participants and runs in each method. The averaged matrix of novel method was shown in Supplementary Fig. S1. Note that the FC matrix was asymmetric due to the method. We then found that these novel FCs were correlated with the seed ROI size. In fact, the correlation coefficient between them was 0.74. Therefore, the effect of ROI size was regressed out from novel FCs before hierarchical clustering. In this study, we averaged the prediction performance from seed A to B with that from B to A for the same ROI pair to assess the novel FC. To transform the similarity measure into the distance measure, both old and novel FCs were converted to one minus the original values. These values were then normalized using min-max normalization. Then, the complete linkage method was applied to both FC patterns. Distance threshold and number of clusters are plotted in Supplementary Fig. S2 (left: traditional method, right: novel method). For the interpretability of the clustering results, the number of clusters should be limited to a relatively small number. In addition, the two numbers should be relatively close in the two methods. The number of repetition of the same distance threshold could be interpreted as a measure of the robustness of the clustering. We determined the number of clusters to be 11 in traditional method and 13 in novel method. To investigate the association between two methods, we searched for the most corresponding pair of clusters between these two clustering results. The correspondence was evaluated by using the algorithm described below. When the number of clusters in the first clustering result is M and the number of clusters in the second clustering result is N, the “overlap ratio matrix” is given as an M × N matrix, where each element consists of the overlap ratio of specific cluster pairs in the two clustering results. When the number of samples x such that x ∈cluster A and x∈cluster B is L, the number of samples in cluster A is M, and the number of samples in cluster B is N, the ratio is expressed as 2 × L / (M + N). When the overlap ratio between clusters A and B is maximized in both the first and second cluster results, we define that clusters A and B correspond. The dendrograms of two clustering results were generated by using the scipy function cluster.hierarchy.dendrogram. Corresponding clusters were visualized in these dendrograms. Moreover, surface maps representing the clustering results were visualized using the Nilearn function plotting.view.surf.

### 2.4 Prediction for individual human traits and behaviors

Next, we compared the predictive performances for human traits and behaviors between the support vector machine (SVM) model built from the traditional FC patterns and the ensemble model built from the SVM models built from both the traditional and novel FC patterns. For this purpose, we used sex, age and 58 behavioral measures across cognition, personality, and emotion that have been mentioned in previous studies (Kong et al., 2019, Li et al., 2019, He et al., 2020) (Supplementary Table 1). Details of the data have been reported in Holmes et al., 2015. In this section, the main processing was performed by customized Matlab R2019b programs.

#### 2.4.1 Prediction by the Model of the single FC pattern

The prediction of the human trait and behavior was conducted as described below for each of traditional and novel FC patterns. FC patterns were used as features. sample size was 995 participants × 4 runs. The FC patterns were standardized using a Z-score transformation for each sample. In the 58 behavior predictions, prior to the training, sex and age effects were regressed out from both FC patterns and behavior measures. In a random 10-fold cross-validation, 9 folds were used to train a SVM model as the training set, and the remaining fold was used to fit the model to the test data. These procedures were performed by using the Matlab fitcsvm function in sex classification problem and the Matlab fitrsvm function in other regression problems. Note that data splitting was performed based on participant level to avoid splitting the same participant’s data into both training and test folds. Predictive values were averaged over 4 runs. Pearson’s correlation coefficient was used as the accuracy metric except sex prediction. In the case of sex prediction, classification scores were averaged over 4 runs. In this case, the predicted label was determined by the sign of the averaged score.

#### 2.4.2 Prediction by the ensemble model of two different FC patterns

We adopted the weighted average approach as the ensemble method. This approach was chosen based on the simple idea that a better predictive performance could be obtained by optimally combining the predictive values provided by two models built on two FC patterns that have different information properties. However, if we were to naively apply this idea to the predicted scores of both FC estimation methods in all participants as calculated as described in the last section, it would lead to the data leakage problem because no independent data would remain for optimal weight estimation. To avoid this shortcoming, we adopted the nested cross-validation (CV) approach. A schematic illustration of this is shown in Fig. 1B. The data were divided into 10 folds for the outer CV loop (Fig 1B, left and top). Specifically, all the data were divided into 9-fold outer training data and a remaining outer test data. The 9-fold outer training data were further divided into 8-fold inner training data and a remaining inner validation data for the inner CV loop (Fig. 1B, right and top). For each inner CV loop, 8 folds were used to train an SVM model, and the remaining fold was used to fit the model to the validation data. This procedure gave us the predicted scores in both traditional and novel methods in an inner validation fold. The details of the model fitting were the same as described in the last section. Next, the actual scores and two predicted scores were standardized using a z-score transformation in an inner validation fold. Then, two weights were computed using the least squares method so that the weighted average of the two predicted scores came closest to the actual score. Specifically, the linear least squares problem was solved with the constraint that each weight was 0 or greater and the sum of two weights was 1, using a built-in Matlab function “lsqlin” (See bottom of Fig. 1B). These procedures yielded 9 weights for each FC estimation method and these weights were then averaged (Fig 1B, center in right side). Importantly, these procedures enabled us to obtain weights and predicted scores in both FC estimation methods for an outer test fold without using the outer test data. Finally, the weighted average of the two predicted scores in single FC estimation method was treated as the ensemble predicted score. (Fig. 1B, center in left side).

The predictive performances of the 3 models, namely old method, novel method, and ensemble method in sex classification and age prediction are reported in Supplementary Fig. S3. Also, the predictive performances of the 3 methods in 58 behavioral measures are reported in Supplementary Fig. S4. We statistically compared these performances between traditional and ensemble methods. We performed a McNemar’s mean test to compare their accuracies in sex classification. We used the correlation coefficient as a measure of prediction accuracy in age and 58 behaviors. To statistically compare two predictive accuracies in each individual trait, a statistical test for the difference between two dependent correlation coefficients in each behavioral measure was performed according to the previous textbook (Glass et al., 1996). A two-sample t-test was conducted to test the difference in prediction accuracy between traditional and ensemble models on 58 behaviors. Furthermore, MSE (Mean Squared Error) was computed and used to test the difference between traditional and ensemble learners in 58 behaviors (Supplementary Fig. S5). A test of the difference between two dependent correlation coefficients used in age prediction was also performed in each behavior.

## Results

### 3.1. Two FC patterns computed by different analysis methods have different information properties

We extracted two types of FCs based on the HCP-MMP atlas which includes cortical regions (Glasser et al., 2016). Traditional FC was estimated by calculating correlation coefficients between averaged time courses of fMRI signals in different ROIs. In contrast, novel FC was estimated by using information of the fine-grained spatial signal patterns in ROI (See Fig. 1A. Materials and Methods section). Matrices of these FCs indicate that these patterns based on different methods have different information contents as shown in Fig. 2A. To explore the difference in detail, we performed hierarchical clustering on these FC patterns. For the interpretability of the clustering, the number of clusters should be limited to a relatively small number. Therefore, we set the number of clusters to be 11 in traditional method and 13 in novel method based on robustness considerations (see Supplementary Fig 2). Furthermore, we found corresponding cluster pairs between two FCs based on the ‘overlap ratio’ (see Materials and Methods section). The Fig. 2B shows the dendrograms of the two clustering results (left, old FC; right novel FC) and the corresponding cluster pairs shown in lines of the middle of the figure. Note that the number in bracket in each cluster represents the number of ROIs. Surface maps of these clustering results are shown in Fig. 2C. Note that corresponding clusters were indicated by the same color on two maps. Furthermore, the line color in the center of the Fig. 2B corresponds to the cluster color in the Fig 2C. While corresponding areas are distributed in right frontal and lateral posterior areas, part of parietal, temporal and medial posterior areas are not corresponding. These results suggest that the FC pattern of the novel method have different information properties than that of the traditional method.

**Fig. 2.**
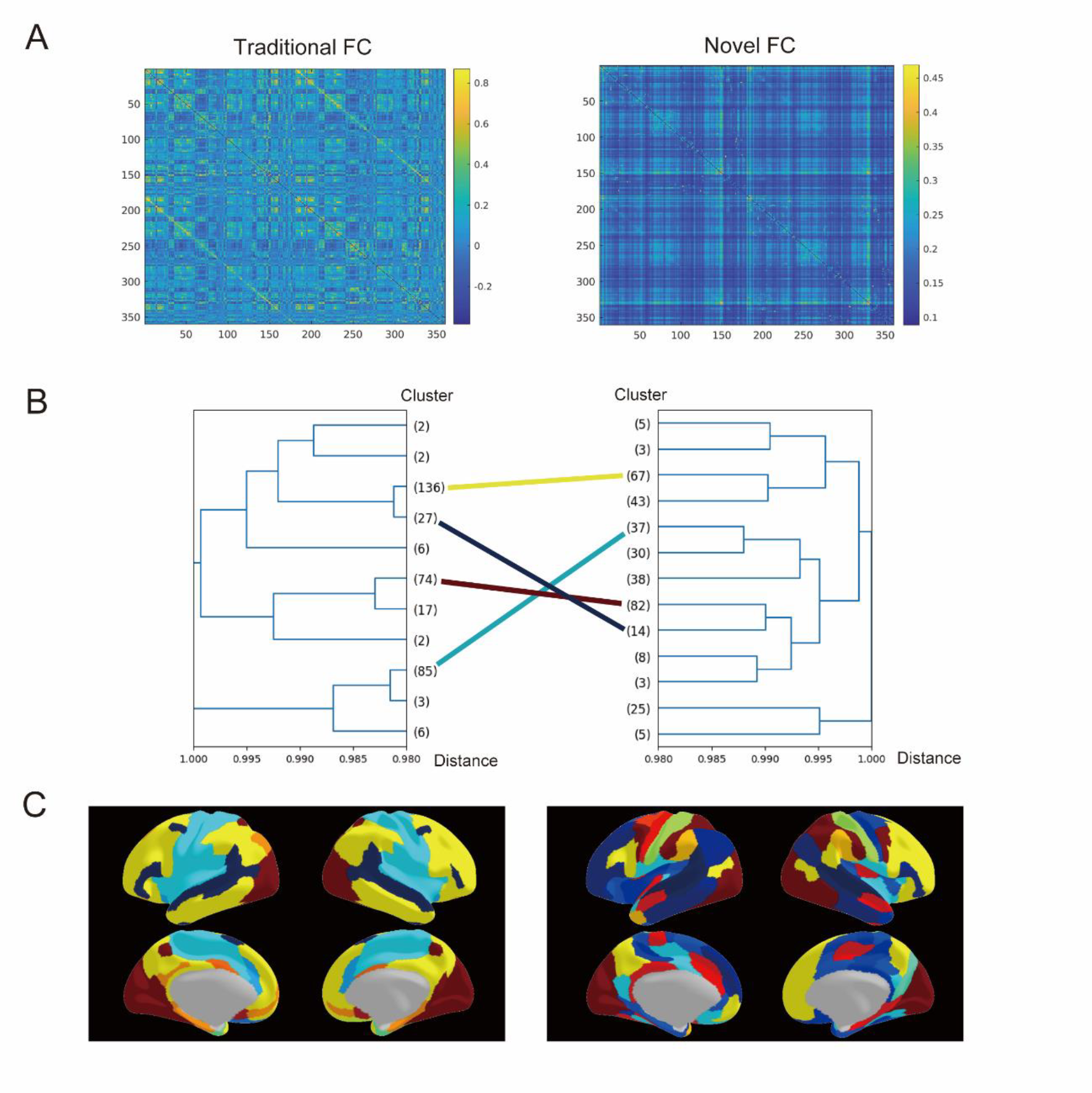
Traditional and novel FC matrices and the hierarchical clustering results. **A:** The FC matrices for traditional (left) and novel (right) methods are depicted. Using Glasser’s scheme, 180 regions of interest (ROIs) are defined for each hemisphere. Traditional FCs are derived by calculating Pearson correlations among time series data across different ROIs, whereas novel FCs result from correlating actual and predicted signals at each target vertex, with predictions based on the local spatial pattern within the ROI. This approach yields two metrics per ROI pair, but for clarity, the matrix displays the average of these metrics for each pair. **B:** Hierarchical clustering results are presented in dendrogram form for both traditional (left) and novel (right) FC patterns. The traditional method identifies 11 clusters, while the novel method finds 13, illustrating the nuanced differences captured by the novel approach. The horizontal axis indicates the ratio of heights, and bracketed values denote the ROI count within each cluster. Lines connecting the dendrograms represent cluster pairs with corresponding methods, with each line’s color matching that of the brain map in Fig. 2C, highlighting the comparative analysis between traditional and novel FC methods. **C:** Brain surface maps illustrate the clustering results for traditional (left) and novel (right) FCs, with corresponding cluster pairs indicated by unified color schemes. This visual representation underscores the distinct connectivity patterns captured by the novel FC analysis compared to the traditional method, offering deeper insights into brain network organization and functional connectivity.

### 3.2. The ensemble model achieves better performance in predicting human traits and behaviors

The hierarchical clustering results suggest possibility that these two types of FCs have different properties of information about brain-behavior relationships. If so, by exploiting such information, it was possible that an ensemble model of these two types of FCs could achieve better prediction performance of individual traits and behaviors than a single model of the traditional FC. To investigate this possibility, we evaluated prediction performance of sex, age, and behavioral measures using 2 FC patterns and an ensemble of them. We built support vector machine models to predict sex, age and 58 behavioral measures by using traditional and novel FC patterns. These 58 measures were commonly used in the previous studies (Kong et al., 2019, Li et al., 2019, He et al., 2020, See Supplementary Table 1). Furthermore, we also built ensemble model of these two methods by using weight average technique. Except sex classification, Pearson correlation coefficient was adopted as a prediction accuracy metric. The results of traditional, novel and ensemble methods are reported in Supplementary Fig. 3 and 4. Furthermore, Fig. 3 shows a comparison of the predictive performances between traditional and ensemble methods with statistical tests with multiple comparisons. While orange color indicates performance of the traditional method, blue color indicates performance of the ensemble method. The sex classification accuracies are shown in Fig. 3A. We tested statistical difference between two classification accuracies by using the mid-P value McNemar’s Test. As a result, there is no difference between them (t2*_(1)_ = 20, p = 0.65). The age prediction accuracies are shown in Fig. 3B. We tested the statistical difference between two dependent correlations by using a classical test technique (Glass et al., 1996). Note that this technique was used in all comparisons of correlations between traditional and ensemble methods in this section. As a result, the ensemble performance is significantly higher than the traditional one (t_(992)_ = 2.89, p = 3.92×10^−3^). Furthermore, we conducted a paired sample t-test of the correlation coefficient in 58 measures between traditional and ensemble methods. As a result, ensemble accuracies are significantly higher than traditional ones (t_(57)_ = 6.94, p = 4.06 × 10^−9^) (Fig. 3C). To confirm the robustness of this result by using another metric, we performed a paired sample t-test of the MSE (Mean Squared Error) in 58 measures between traditional and ensemble methods. As a result, the ensemble accuracies of the ensemble methods are significantly lower than the traditional ones (t_(57)_ = 4.64, p = 2.11×10^−5^) (Supplementary Fig. S5). To investigate these effects in each behavior, we tested the statistical difference between two dependent correlations in each measurement in the same way as for age prediction. After correction for multiple comparison, ensemble performances are significantly higher than the traditional ones in one measure with Bonferroni correction and with 3 measures in Holm correction. There is no measure in which traditional method whose performance is significantly higher than the ensemble one. To show in which measure the ensemble method achieves higher performance, the representative top 10 measures which are sorted in the rank of the t-values and plotted in Fig. 3D. To visualize the effect of the ensemble method, the scatter plot matrix of the delay discounting measure is shown in Fig.4. Interestingly, the correlation coefficient of the predicted scores of traditional and novel methods were relatively weak as 0.50. These results indicated that two SVM models exploited different information from traditional and novel FCs. Therefore, we concluded that FC extracted by using the local fine-grained spatial activation patterns is beneficial to improve the predictive accuracy of the individual traits and behaviors.

**Fig. 3.**
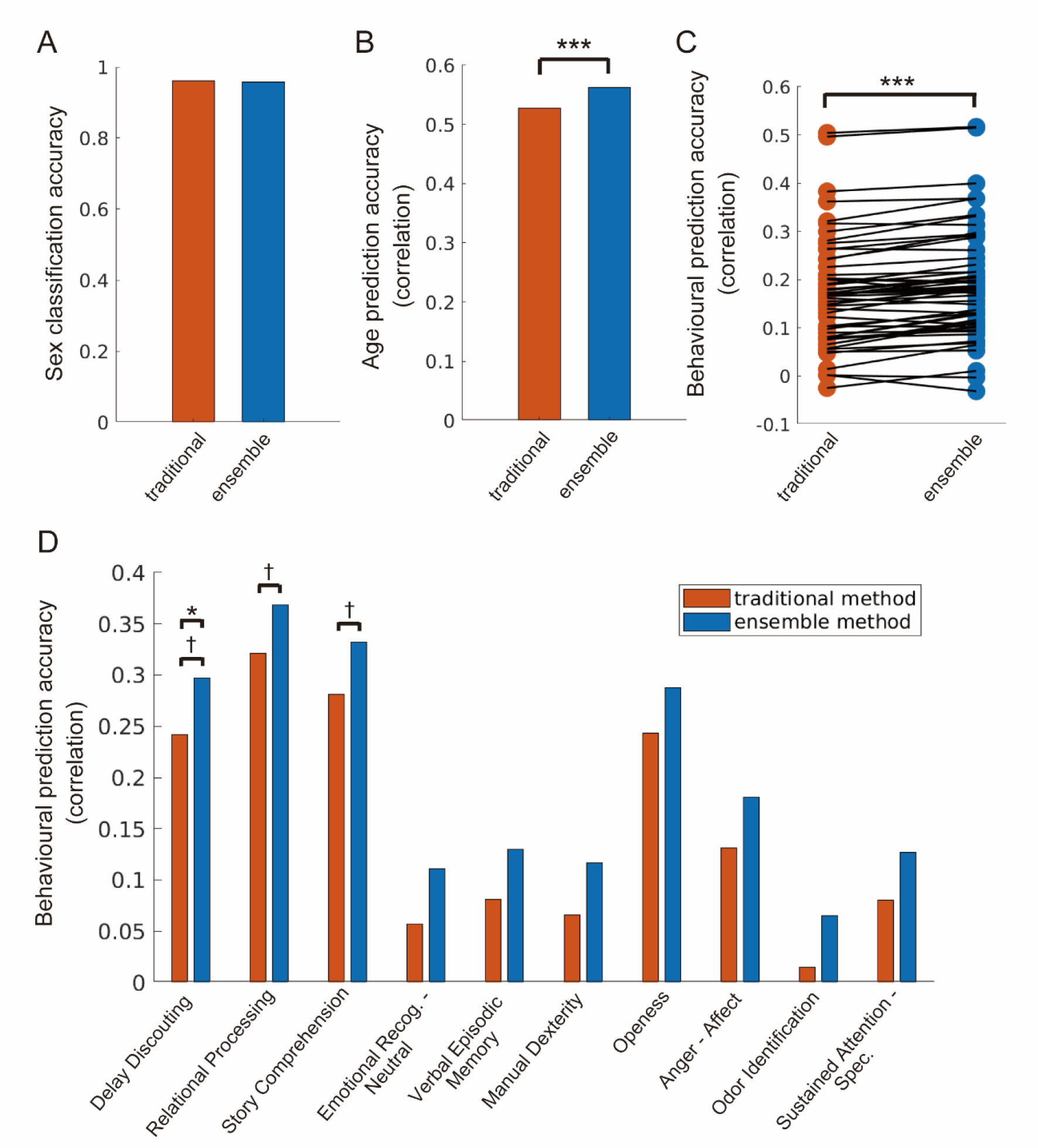
Comparison of predictive performances in human traits and behaviors between traditional and ensemble models. Note that Peason correlation coefficient is used as a predictive accuracy measure without sex classification. **A**: Sex prediction. There is no significant difference between two models. **B**: Age prediction. The ensemble predictive performance is significantly superior to traditional one. **C**: Prediction for the 58 behavioral measures. The ensemble predictive performance of is significantly superior to traditional one in all measures. **D**: Top 10 Predictive performances sorted in the rank of t-value representing the degree of the difference between two models. * p < 0.05, ** p < 0.01, *** p < 0.001 (Multiple comparison with Bonfferoni method), † p < 0.05, †† p < 0.01, ††† p < 0.001 (Multiple comparison with Holm method).

**Fig. 4.**
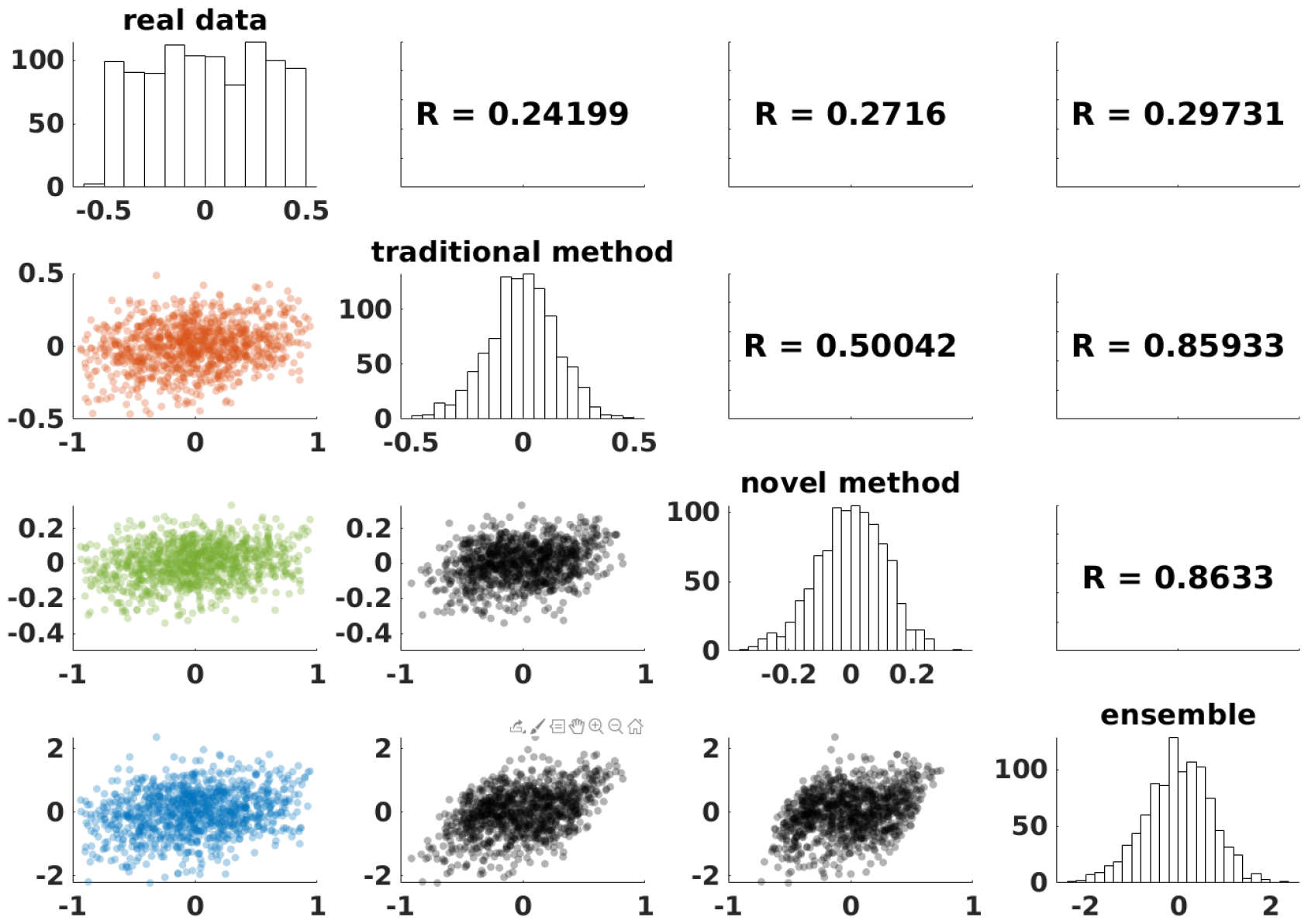
Scatter plot matrix among real data, predicted data by traditional, novel and ensemble methods in delay discounting. Diagonal elements contain histogram of each data. Elements in upper triangle contain Pearson correlation coefficient of each data pair. Elements in lower triangle contain scatter plot in each data pair.

We develop the novel analysis method to uncover functional connectivity by using fine-grained local spatial patterns of the neural activations. Hierarchical clustering shows that clustering result of the novel FC has different grouping patterns in the whole brain compared to that of the traditional FC. Ensemble model of two different FCs achieves better predictive performance in human traits and behaviors than the single model of the traditional FC. Interestingly, similarity of predicted values of traditional and novel methods is relatively weak. In conclusion, our novel method provides different connectivity patterns revealed by using fine-grained local spatial patterns of neural activation than the traditional method. These patterns contain rich and previously unknown information about human traits and behaviors.

## Discussion

In this study, we introduced a novel approach to analyze resting-state functional connectivity by focusing on spatial patterns of local neural activity, information that may be overlooked by conventional methods. Hierarchical clustering of RSFC using both traditional and novel approaches revealed distinct association patterns, with similarities noted in the visual and right frontal cortices. However, divergent clustering results emerged in other brain regions. Additionally, we evaluated the predictive accuracy for sex, age, and 58 behavioral measures using models based on traditional RSFC alone versus an ensemble model combining both traditional and novel RSFC methods. The ensemble model demonstrated significantly improved predictive accuracy for age and various behavioral measures. These findings indicate that RSFC analysis, which incorporates local spatial patterns, captures uniquely valuable information beyond what is obtained through traditional methods. This enriched information substantially enhances the prediction of human traits and behaviors, underscoring the potential of integrating traditional and novel RSFC analyses in cognitive neuroscience.

### 4.1. Novel type of resting-state functional connectivity reflecting local spatial activation patterns

Multi-voxel pattern analysis (MVPA) is an analysis technique that has been developed by actively using information encoded in multiple local voxels / vertices (Haynes, 2015). This technique exploits information that would be lost if the information of individual voxels / vertices were treated separately or averaged (Anzellotti and Coutanche, 2018). MVPA allows for decoding multi-sensory stimuli such as of the visual (Haxby et al., 2001, Kamitani and Tong, 2005), auditory (Hoefle et al., 2018), and gustatory (Chikazoe et al., 2019) modalities. Furthermore, the technique showed that local spatial patterns reflect abstract and complex neural representations such as subjective memory (Rissman et al., 2010), estimation of others’ emotions (Skerry et Saxe., 2015), and cognitive maps (Schuck et al., 2016). However, it has in principle been developed as an analysis method specific to neural representation during explicit task because of its basic assumption that local neural activation patterns index the structure of specific current mental representations and processes (Peelen et Downing, 2023).

In recent studies, human resting-state functional connectivity measured with fMRI have been widely used to construct models to predict human traits and behaviors such as sex (Fan et al., 2020, Zhang et al., 2018), age (Geerligs et al., 2015, Varangis et al., 2019, Kim et al., 2022), personality tendencies (Hsu et al., 2018,Gentili et al., 2017), intelligence (Fan et al., 2020,Hearne et al., 2016), performance on psychological tasks (Liégeois., 2019, Li et al. He et al., 2020), and classification of neuropsychiatric disorders (Yamashita et al., 2020, Yahata et al., 2016, Rashid et al., 2016). Although such studies have overwhelmingly used correlation coefficients between time-series signals averaged over divided brain units such as ROIs, parcels or networks as connectivity metrics (Bijsterbosch et al., 2020), MVPA results suggest that the local spatial patterns of the human fMRI signals contain rich information about human traits and behaviors. In addition, there has been much development of analysis methods focusing on fluctuations in time-series patterns, collectively called dynamic functional connectivity (dFC) (Preti et al., 2017), but in contrast, few have extracted information on spatial patterns. The improved predictive performance of our novel method on human traits and behaviors in this study suggests that local fine-grained spatial patterns of fMRI signals have rich information about human traits and behaviors, not only during explicit task, but also at rest.

### 4.2. Potential Applications as Biomarkers of Neuropsychiatric Disorders

It has been frequently noted that the diagnostic criteria for neuropsychiatric disorders largely rely on self-reported internal experiences and observed behaviors of patients, due to the absence of reliable biological markers (Kapur et al., 2012; Parkers et al., 2020; Woo et al., 2017). Consequently, the identification and validation of biological biomarkers for neuropsychiatric conditions, including depression, schizophrenia, and autism spectrum disorders (ASD), represent critical challenges for contemporary psychiatry. Numerous efforts are underway to develop classifiers capable of distinguishing neuropsychiatric disorders by leveraging advanced machine learning techniques applied to resting-state functional connectivity (RSFC) data, as captured through functional magnetic resonance imaging (fMRI). For example, Drysdale et al. showed that depression can be clustered into four biotypes based on large-scale fMRI data, which enabled us to predict the therapeutic effect of the transcranial magnetic therapy (Drysdale et al., 2017). Yahata et al. also developed a classifier algorithm using a small number of resting-state functional connectivity that provided the generalized predictive performance to classify ASD and neurotypical population across different international cohorts (Yahata et al., 2016). Lei et al. showed that classifier based on integrating data from structural and functional images achieved better classification performance than that based on a single data (Lei et al., 2019). These results indicate that neuroimaging methods have potential not only for elucidating the pathophysiology of neuropsychiatric disorders but also for developing the clinical applications. However, most classifiers for neuropsychiatric disorders utilizing resting-state functional connectivity rely on averaged time-series correlations within regions of interest (ROIs) as a measure of connectivity. This approach could overlook crucial information encoded in local spatial patterns, which are significant for understanding human behavior and psychopathology.

We have devised a method to analyze resting-state functional connectivity that captures the spatial patterns of local neural activity often overlooked by conventional approaches. The novel method for calculating resting-state functional connectivity revealed connectivity patterns distinct from those identified by the conventional approach, which relies on averaged time-series correlations within ROIs. We assessed the predictive performance of models based on traditional connectivity against those derived from an ensemble approach, incorporating both traditional and novel functional connectivity methods. The ensemble model demonstrated superior performance compared to the traditional model alone. These findings indicate that our novel approach to functional connectivity uncovers detailed information pertinent to human traits and behaviors. Given its potential benefits, this novel method could significantly contribute to developing classifiers for neuropsychiatric disorders, marking a promising direction for future research.

## Data and code availability

All data used in the present study publicly available data of resting-state fMRI distributed by the Human Connectome Project (HCP; http://humanconnectomeproject.org/). Codes for main analyses are available in GitHub at https://github.com/takaakiyoshim/Novel_Method_For_RSFC.

## Author Contributions

Conceptualization, T.Y. and J.C. Methodology, T.Y. and J.C. Investigation, T.Y., K.T. and J.C. Writing – Original Draft, T.Y. Writing – Review & Editing, J.C. Supervision, J.C. Funding Acquisition, J.C.

## Declaration of Competing Interests

No, I declare the authors have no competing interests as defined by Imaging Neuroscience, or other interests that might be perceived to influence the interpretation of the article.

## Acknowledgements

Computational resources were provided by the Data Integration and Analysis Facility, National Institute for Basic Biology, and Research Center for Computational Science.

## Notes

### Competing Interest Statement

The authors have declared no competing interest.

## Reference

Anzellotti, S., Coutanche, M.N., 2018. Beyond Functional Connectivity: Investigating Networks of Multivariate Representations. Trends Cogn. Sci. 22, 258–269. doi: 10.1016/j.tics.2017.12.002

Biswal, B., Yetkin, F.Z., Haughton, V.M., Hyde, J.S., 1995. Functional connectivity in the motor cortex of resting human brain using echo-planar MRI. Magn. Reson. Med. 34, 537–541. doi: 10.1002/mrm.1910340409

Chikazoe, J., Lee, D.H., Kriegeskorte, N., Anderson, A.K., 2019. Distinct representations of basic taste qualities in human gustatory cortex. Nat. Commun. 10, 1048. doi: 10.1038/s41467-019-08857-z

Drysdale, A.T., Grosenick, L., Downar, J., Dunlop, K., Mansouri, F., Meng, Y., Fetcho, R.N., Zebley, B., Oathes, D.J., Etkin, A., Schatzberg, A.F., Sudheimer, K., Keller, J., Mayberg, H.S., Gunning, F.M., Alexopoulos, G.S., Fox, M.D., Pascual-Leone, A., Voss, H.U., Casey, B.J., Dubin, M.J., Liston, C., 2017. Resting-state connectivity biomarkers define neurophysiological subtypes of depression. Nat. Med. 23, 28–38. doi: 10.1038/nm.4246

Fan, L., Su, J., Qin, J., Hu, D., Shen, H. 2020. A Deep Network Model on Dynamic Functional Connectivity With Applications to Gender Classification and Intelligence Prediction. Front. Neurosci. 14, 881. doi: 10.3389/fnins.2020.00881

Geerligs, L., Renken, R.J., Saliasi, E., Maurits, N.M., Lorist, M.M., 2015. A Brain-Wide Study of Age-Related Changes in Functional Connectivity. Cereb. Cortex 25, 1987–1999. doi: 10.1093/cercor/bhu012

Gentili, C., Cristea, I.A., Ricciardi, E., Vanello, N., Popita, C., David, D., Pietrini, P., 2017. Not in one metric: Neuroticism modulates different resting state metrics within distinctive brain regions. Behav. Brain Res. 327, 34–43. doi: 10.1016/j.bbr.2017.03.031

Glass, G.V., & Hopkins, K.D. (1996). Statistical Methods in Education and Psychology (3rd Ed.). Boston: Allyn & Bacon. (Chapter 14)

Glasser, M.F., Sotiropoulos, S.N., Wilson, J.A., Coalson, T.S., Fischl, B., Andersson, J.L., Xu, J., Jbabdi, S., Webster, M., Polimeni, J.R., et al., 2013. The minimal preprocessing pipelines for the human connectome project. Neuroimage 80, 105–124. doi: 10.1016/j.neuroimage.2013.04.127.

Glasser, M.F., Coalson, T.S., Robinson, E.C., Hacker, C.D., Harwell, J., Yacoub, E., Ugurbil, K., Andersson, J., Beckmann, C.F., Jenkinson, M., Smith, S.M., Van Essen, D.C., 2016. A multi-modal parcellation of human cerebral cortex. Nature 536, 171–178. doi: 10.1038/nature18933

Haynes, J.-D., Rees, G., 2005. Predicting the orientation of invisible stimuli from activity in human primary visual cortex. Nat. Neurosci. 8, 686–691. doi: 10.1038/nn1445

Haynes, J.-D., 2015. A Primer on Pattern-Based Approaches to fMRI: Principles, Pitfalls, and Perspectives. Neuron 87, 257–270. doi: 10.1016/j.neuron.2015.05.025

He, T., Kong, R., Holmes, A.J., Nguyen, M., Sabuncu, M.R., Eickhoff, S.B., Bzdok, D., Feng, J., Yeo, B.T.T., 2020. Deep neural networks and kernel regression achieve comparable accuracies for functional connectivity prediction of behavior and demographics. Neuroimage 206, 116276. doi:10.1016/j.neuroimage.2019.116276

Hearne, L.J., Mattingley, J.B., Cocchi, L., 2016. Functional brain networks related to individual differences in human intelligence at rest. Sci. Rep. 6, 32328. doi: 10.1038/srep32328

Heinzle, J., Kahnt, T., Haynes, J.-D., 2011. Topographically specific functional connectivity between visual field maps in the human brain. Neuroimage 56, 1426–1436. doi: 10.1016/j.neuroimage.2011.02.077

Hoefle, S., Engel, A., Basilio, R., Alluri, V., Toiviainen, P., Cagy, M., Moll, J., 2018. Identifying musical pieces from fMRI data using encoding and decoding models. Sci. Rep. 8, 2266. doi: /10.1038/s41598-018-20732-3

Hsu, W.-T., Rosenberg, M.D., Scheinost, D., Constable, R.T., Chun, M.M., 2018. Resting-state functional connectivity predicts neuroticism and extraversion in novel individuals. Soc. Cogn. Affect. Neurosci. 13, 224–232. doi: 10.1093/scan/nsy002

Kalmady, S.V., Greiner, R., Agrawal, R., Shivakumar, V., Narayanaswamy, J.C., Brown, M.R.G., Greenshaw, A.J., Dursun, S.M., Venkatasubramanian, G., 2019. Towards artificial intelligence in mental health by improving schizophrenia prediction with multiple brain parcellation ensemble-learning. NPJ Schizophr 5, 2. doi: 10.1038/s41537-018-0070-8

Kamitani, Y., Tong, F., 2005. Decoding the visual and subjective contents of the human brain. Nat. Neurosci. 8, 679–685. doi: 10.1038/nn1444

Kapur, S., Phillips, A.G., Insel, T.R., 2012. Why has it taken so long for biological psychiatry to develop clinical tests and what to do about it? Mol. Psychiatry 17, 1174–1179. doi: 10.1038/mp.2012.105

Khosla, M., Jamison, K., Kuceyeski, A., Sabuncu, M.R., 2019. Ensemble learning with 3D convolutional neural networks for functional connectome-based prediction. Neuroimage 199, 651– 662. doi: 10.1016/j.neuroimage.2019.06.012

Kim, E., Kim, S., Kim, Y., Cha, H., Lee, H.J., Lee, T., Chang, Y., 2022. Connectome-based predictive models using resting-state fMRI for studying brain aging. Exp. Brain Res. 240, 2389–2400. doi: 10.1007/s00221-022-06430-7

Lei, D., Pinaya, W.H.L., Young, J., van Amelsvoort, T., Marcelis, M., Donohoe, G., Mothersill, D.O., Corvin, A., Vieira, S., Huang, X., Lui, S., Scarpazza, C., Arango, C., Bullmore, E., Gong, Q., McGuire, P., Mechelli, A., 2020. Integrating machining learning and multimodal neuroimaging to detect schizophrenia at the level of the individual. Hum. Brain Mapp. 41, 1119–1135. doi: 10.1002/hbm.24863

Li, J., Kong, R., Liégeois, R., Orban, C., Tan, Y., Sun, N., Holmes, A.J., Sabuncu, M.R., Ge, T., Yeo, B.T.T., 2019. Global signal regression strengthens association between resting-state functional connectivity and behavior. Neuroimage 196, 126–141. doi: 10.1016/j.neuroimage.2019.04.016

Liégeois, R., Li, J., Kong, R., Orban, C., Van De Ville, D., Ge, T., Sabuncu, M.R., Yeo, B.T.T., 2019. Resting brain dynamics at different timescales capture distinct aspects of human behavior. Nat. Commun. 10, 2317. doi: 10.1038/s41467-019-10317-7

Parkes, L., Satterthwaite, T.D., Bassett, D.S., 2020. Towards precise resting-state fMRI biomarkers in psychiatry: synthesizing developments in transdiagnostic research, dimensional models of psychopathology, and normative neurodevelopment. Curr. Opin. Neurobiol. 65, 120–128. doi: 10.1016/j.conb.2020.10.016

Peelen, M.V. & Downing, P.E, 2023. Testing cognitive theories with multivariate pattern analysis of neuroimaging data. Nat Hum Behav 7, 1430–1441. doi 10.1038/s41562-023-01680-z

Preti, M.G., Bolton, T.A., Van De Ville, D., 2017. The dynamic functional connectome: State-of-the-art and perspectives. Neuroimage 160, 41–54. doi: 10.1016/j.neuroimage.2016.12.061

Qin, J., Chen, S.-G., Hu, D., Zeng, L.-L., Fan, Y.-M., Chen, X.-P., Shen, H., 2015. Predicting individual brain maturity using dynamic functional connectivity. Front. Hum. Neurosci. 9, 418. doi: 10.3389/fnhum.2015.00418

Rashid, B., Arbabshirani, M.R., Damaraju, E., Cetin, M.S., Miller, R., Pearlson, G.D., Calhoun, V.D., 2016. Classification of schizophrenia and bipolar patients using static and dynamic resting-state fMRI brain connectivity. Neuroimage 134, 645–657. doi: 10.1016/j.neuroimage.2016.04.051

Rissman, J., Greely, H.T., Wagner, A.D., 2010. Detecting individual memories through the neural decoding of memory states and past experience. Proc. Natl. Acad. Sci. U. S. A. 107, 9849–9854. doi: 10.1073/pnas.1001028107

Schuck, N.W., Cai, M.B., Wilson, R.C., Niv, Y., 2016. Human Orbitofrontal Cortex Represents a Cognitive Map of State Space. Neuron 91, 1402–1412. doi: 10.1016/j.neuron.2016.08.019

Skerry, A.E., Saxe, R., 2015. Neural representations of emotion are organized around abstract event features. Curr. Biol. 25, 1945–1954.

Van Essen, D.C., Smith, S.M., Barch, D.M., Behrens, T.E., Yacoub, E., Ugurbil, K., Consortium, W.M.H., 2013. The WU-Minn human connectome project: an overview. Neuroimage 80, 62–79. doi: 10.1016/j.neuroimage.2013.05.041.

Varangis, E., Habeck, C.G., Razlighi, Q.R., Stern, Y., 2019. The Effect of Aging on Resting State Connectivity of Predefined Networks in the Brain. Front. Aging Neurosci. 11, 234. doi: 10.3389/fnagi.2019.00234

Woo, C.-W., Chang, L.J., Lindquist, M.A., Wager, T.D., 2017. Building better biomarkers: brain models in translational neuroimaging. Nat. Neurosci. 20, 365–377. doi: 10.1038/nn.4478

Yahata, N., Morimoto, J., Hashimoto, R. et al. A small number of abnormal brain connections predicts adult autism spectrum disorder. Nat Commun 7, 11254 (2016). doi: 10.1038/ncomms11254

Yamashita, A., Sakai, Y., Yamada, T., Yahata, N., Kunimatsu, A., Okada, N., Itahashi, T., Hashimoto, R., Mizuta, H., Ichikawa, N., Takamura, M., Okada, G., Yamagata, H., Harada, K., Matsuo, K., Tanaka, S.C., Kawato, M., Kasai, K., Kato, N., Takahashi, H., Okamoto, Y., Yamashita, O., Imamizu, H., 2020. Generalizable brain network markers of major depressive disorder across multiple imaging sites. PLoS Biol. 18, e3000966. doi: 10.1371/journal.pbio.3000966

Zhang, C., Dougherty, C.C., Baum, S.A., White, T., Michael, A.M., 2018. Functional connectivity predicts gender: Evidence for gender differences in resting brain connectivity. Hum. Brain Mapp. 39, 1765–1776. doi: 10.1002/hbm.23950

Zhou Z H. Ensemble Methods: Foundations and Algorithms. Chapman and Hall/CRC, 2012

