## Supplementary_Materials for "Enhancing Prediction of Human Traits and Behaviors through Ensemble Learning of Traditional and Novel Resting-State fMRI Connectivity Analyses"

|  | **Description** | **Field name** |
| --- | --- | --- |
| 1 | Visual Episodic Memory | PicSeq_Unadj |
| 2 | Cognitive Flexibility (DCCS) | CardSort_Unadj |
| 3 | Inhibition (Flanker Task) | Flanker_Unadj |
| 4 | Fluid Intelligence (PMAT) | PMAT24_A_CR |
| 5 | Vocabulary (Pronunciation) | ReadEng_Unadj |
| 6 | Vocabulary (Picture Matching) | PicVocab_Unadj |
| 7 | Processing Speed | ProcSpeed_Unadj |
| 8 | Delay Discounting | DDic_AUC_40K |
| 9 | Spatial Orientation | VSPLOT_TC |
| 10 | Sustained Attention – Sens. | SCPT_SEN |
| 11 | Sustained Attention – Spec. | SCPT_SPEC |
| 12 | Verbal Episodic Memory | IWRD_TOT |
| 13 | Working Memory (List Sorting) | ListSort_Unadj |
| 14 | Cognitive Status (MMSE) | MMSE_Score |
| 15 | Sleep Quality (PSQI) | PSQI_Score |
| 16 | Walking Endurance | Endurance_Unadj |
| 17 | Walking Speed | GaitSpeed_Unadj |
| 18 | Manual Dexterity | Dexterity_Unadj |
| 19 | Grip Strength | Strength_Unadj |
| 20 | Odor Identification | Odor_Unadj |
| 21 | Pain Interference Survey | PainInterf_Tscore |
| 22 | Taste Intensity | Taste_Unadj |
| 23 | Contrast Sensitivity | Mars_Final |
| 24 | Emotional Face Matching | Emotion_Task_Face_Acc |
| 25 | Arithmetic | Language_Task_Math_Avg_Difficulty_Level |
| 26 | Story Comprehension | Language_Task_Story_Avg_Difficulty_Level |
| 27 | Relational Processing | Relational_Task_Acc |
| 28 | Social Cognition – Random | Social_Task_Perc_Random |
| 29 | Social Cognition – Interaction | Social_Task_Perc_TOM |

**Table S1.** Lookup table showing the original HCP variable names with the corresponding descriptive labels used in this study.

|  | **Description** | **Field name** |
| --- | --- | --- |
| 30 | Working Memory (N-back) | WM_Task_Acc |
| 31 | Agreeableness (NEO) | NEOFAC_A |
| 32 | Openness (NEO) | NEOFAC_O |
| 33 | Conscientiousness (NEO) | NEOFAC_C |
| 34 | Neuroticism (NEO) | NEOFAC_N |
| 35 | Extraversion (NEO) | NEOFAC_E |
| 36 | Emot. Recog. – Total | ER40_CR |
| 37 | Emot. Recog. – Angry | ER40ANG |
| 38 | Emot. Recog. – Fear | ER40FEAR |
| 39 | Emot. Recog. – Happy | ER40HAP |
| 40 | Emot. Recog. - Neutral | ER40NOE |
| 41 | Emot. Recog. – Sad | ER40SAD |
| 42 | Anger – Affect | AngAffect_Unadj |
| 43 | Anger – Hostility | AngHostil_Unadj |
| 44 | Anger – Aggression | AngAggr_Unadj |
| 45 | Fear – Affect | FearAffect_Unadj |
| 46 | Fear – Somatic Arousal | FearSomat_Unadj |
| 47 | Sadness | Sadness_Unadj |
| 48 | Life Satisfaction | LifeSatisf_Unadj |
| 49 | Meaning & Purpose | MeanPurp_Unadj |
| 50 | Positive Affect | PosAffect_Unadj |
| 51 | Friendship | Friendship_Unadj |
| 52 | Loneliness | Loneliness_Unadj |
| 53 | Perceived Hostility | PercHostil_Unadj |
| 54 | Perceived Rejection | PercReject_Unadj |
| 55 | Emotional Support | EmotSupp_Unadj |
| 56 | Instrument Support | InstruSupp_Unadj |
| 57 | Perceived Stress | PercStress_Unadj |
| 58 | Self-Efficacy | SelfEff_Unadj |

**Table S1 (cont.).** Lookup table showing the original HCP variable names with the corresponding descriptive labels used in this study.

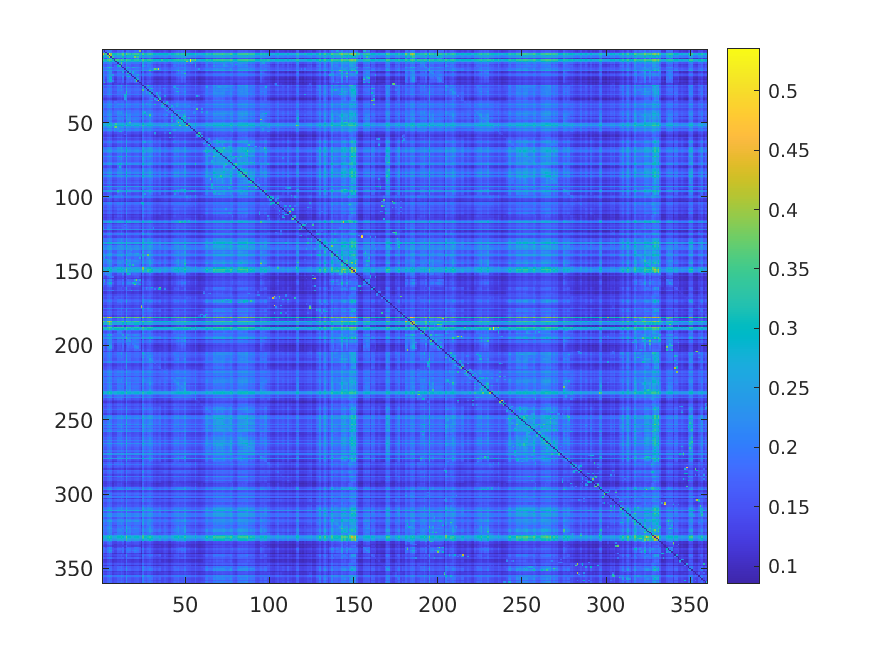

**Figure S1.** The FC matrix in novel method. Note that the matrix is not symmetrical.

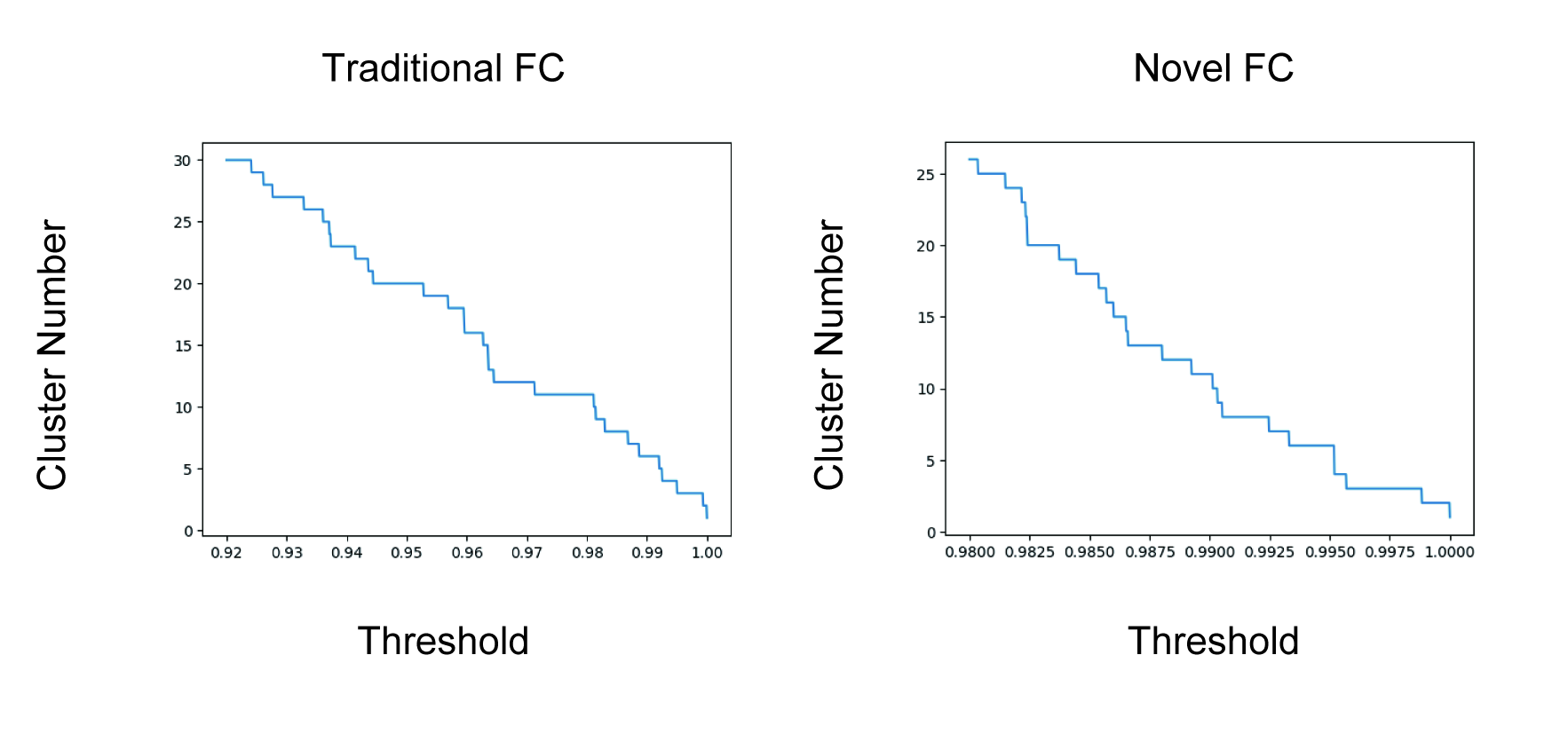

**Figure S2.** Distance threshold and number of clusters of the hierarchical clustering in traditional and novel FC.

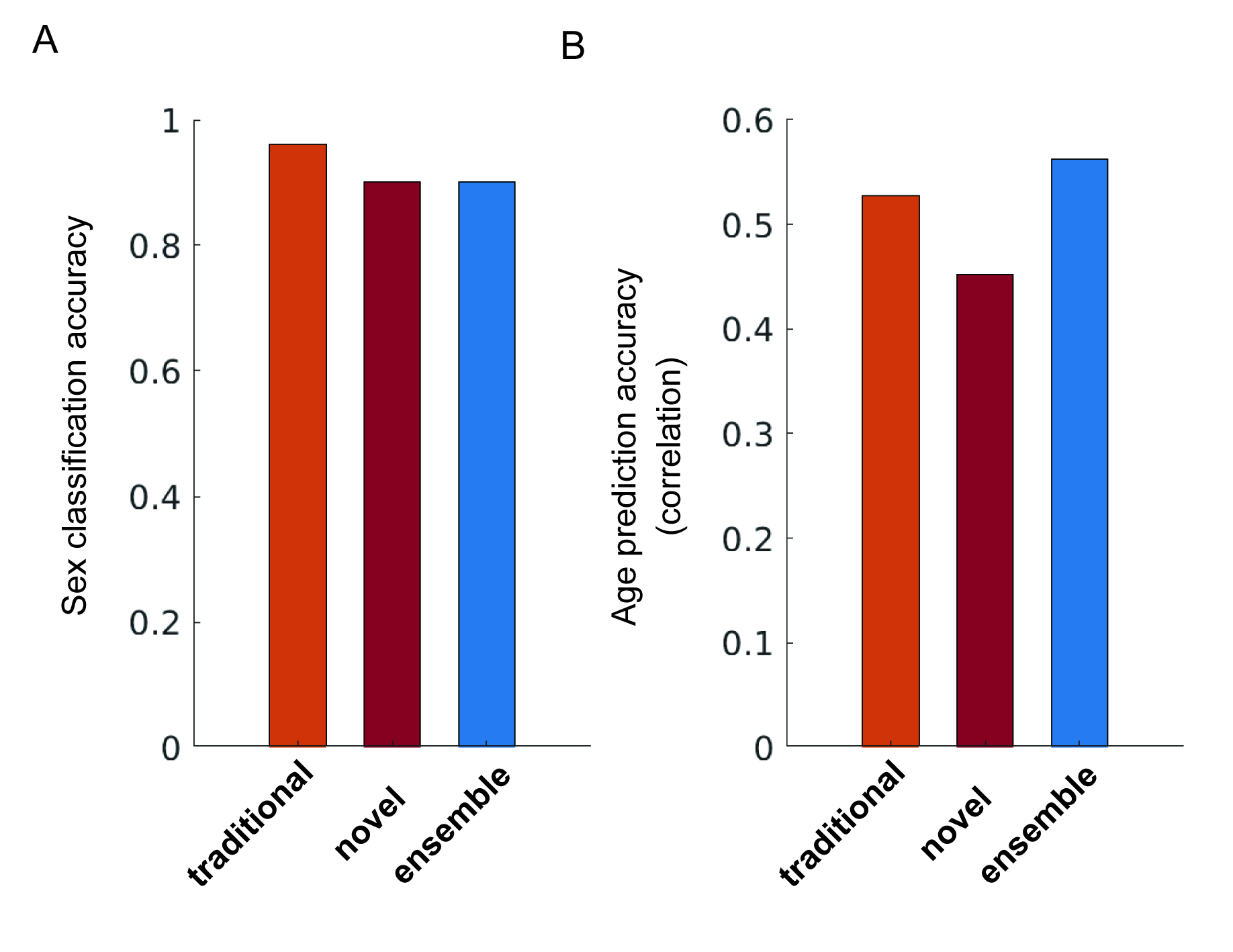

**Figure S3.** Predictive performance of 3 methods in sex classification and age prediction.

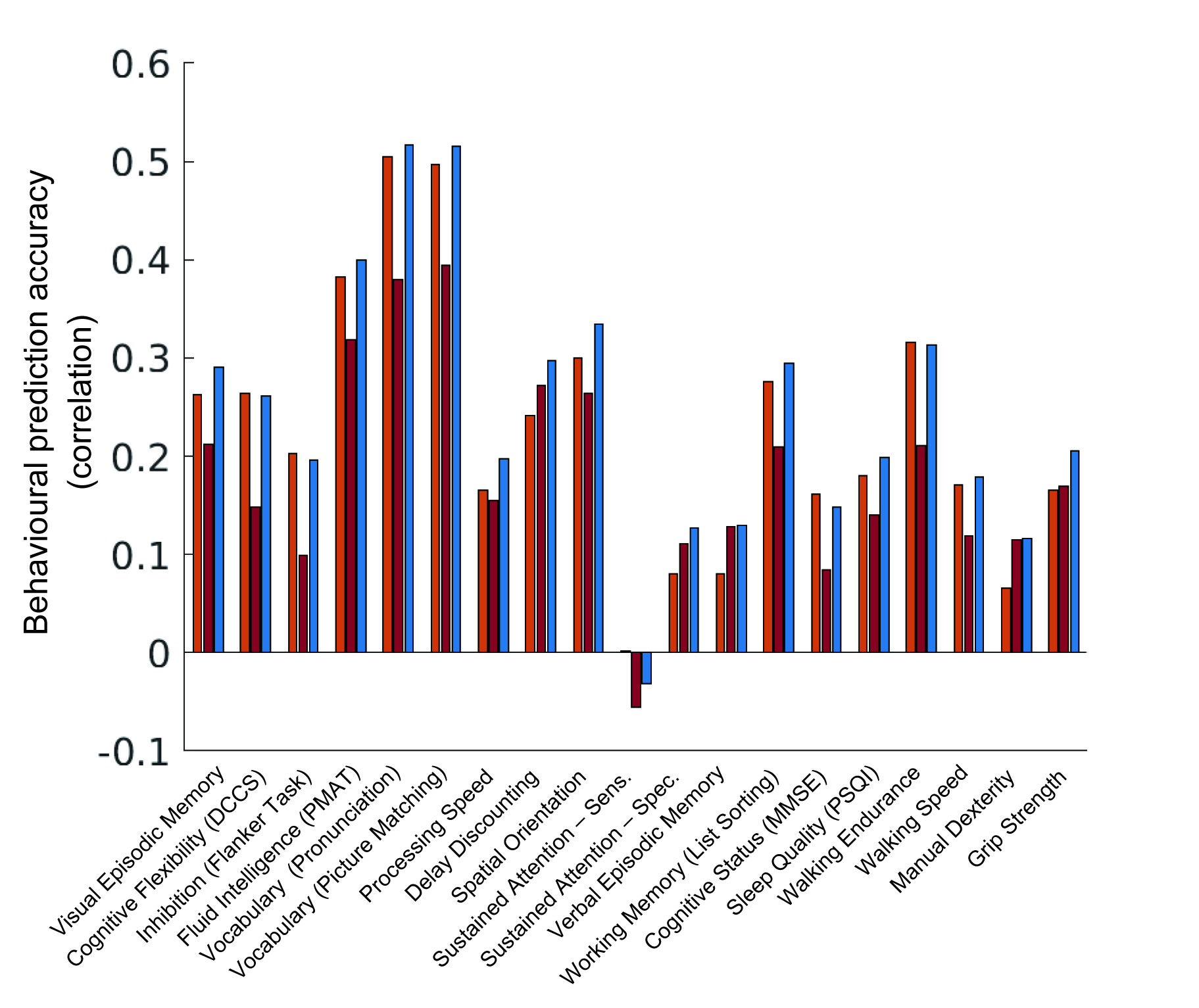

**Figure S4** Predictive performance of 3 methods in individual traits and behaviors. (List No. 1 to No. 19)

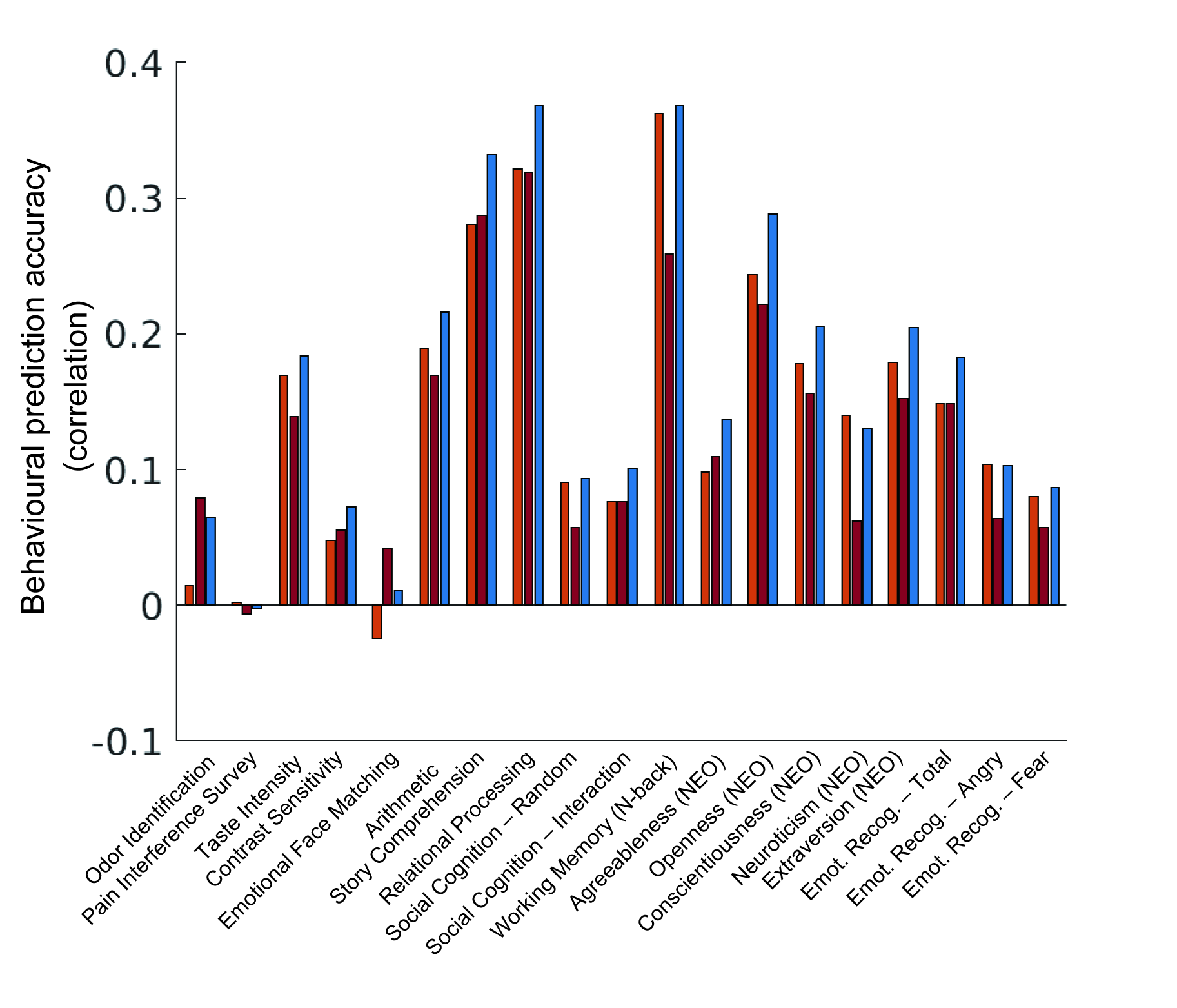

**Figure S4 (cont.).** Predictive performance of 3 methods in individual traits and behaviors. (List No. 20 to No. 38)

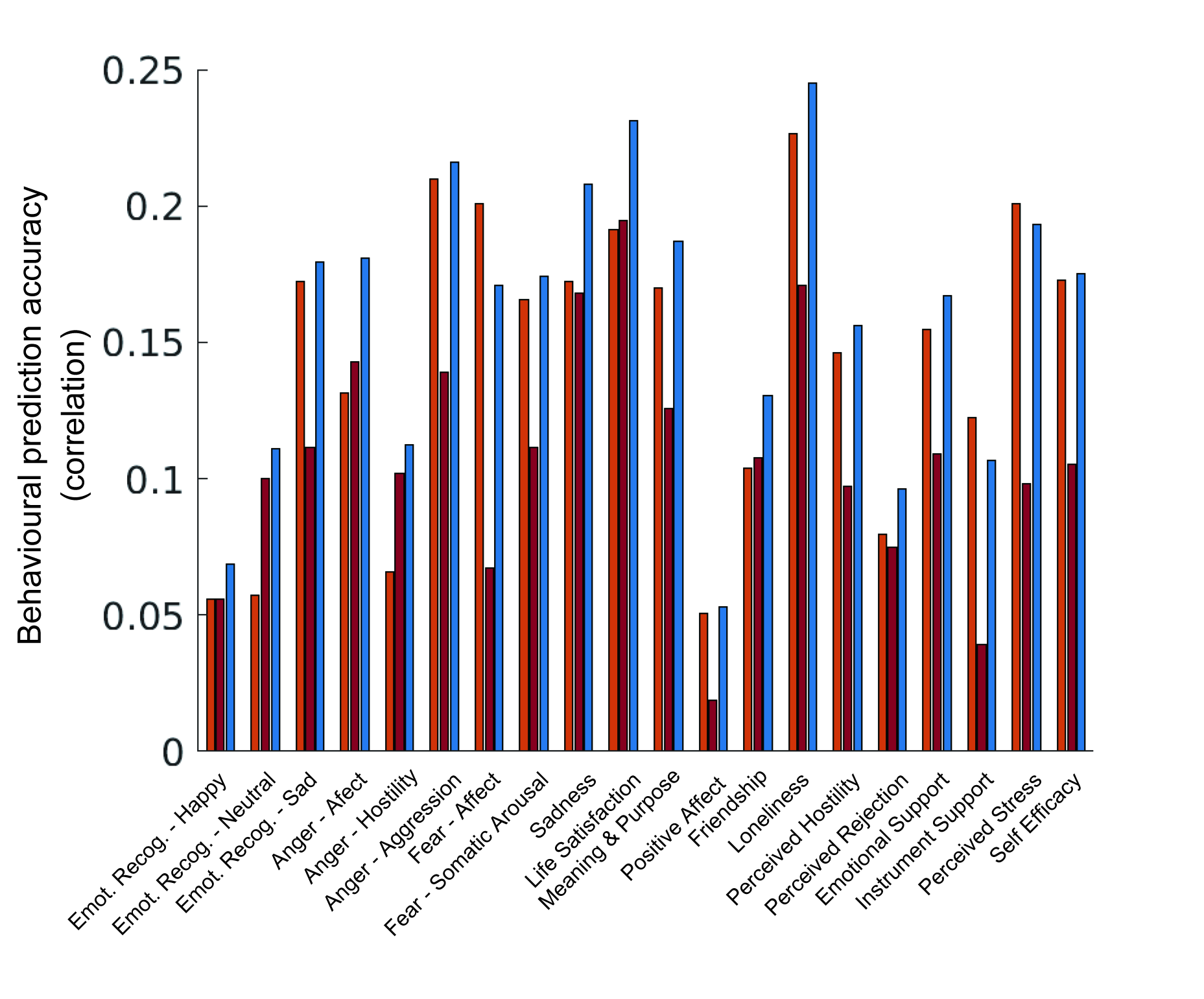

**Figure S4 (cont.).** Predictive performance of 3 methods in individual traits and behaviors. (List No. 39 to No. 58)

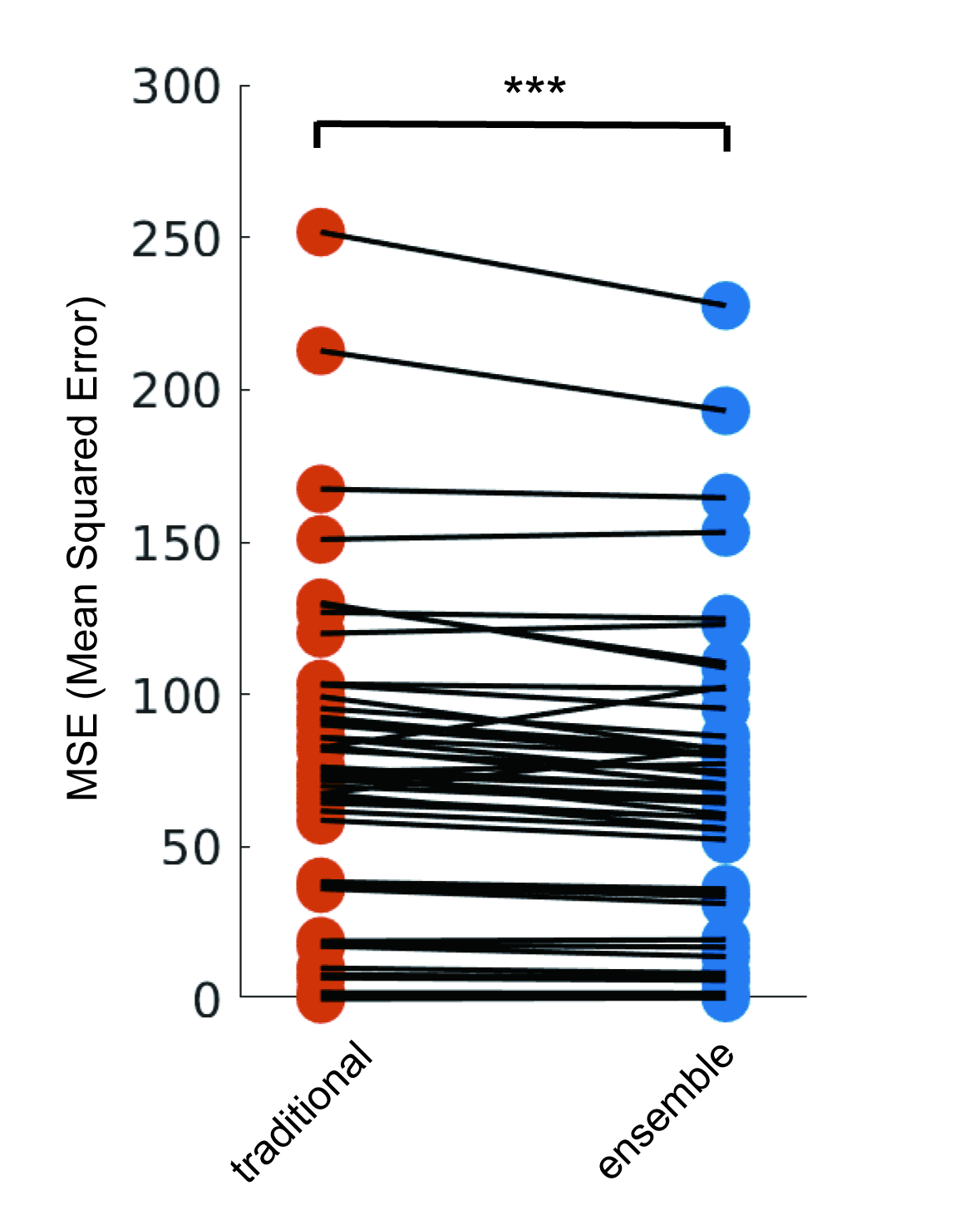

**Figure S5.** Comparison between predictive performances (MSE: Mean Squared Error) of traditional and ensemble methods. * p < 0.05, ** p < 0.01, *** p < 0.001.
